# Parameter inference in dynamical systems with co-dimension 1 bifurcations

**DOI:** 10.1101/623413

**Authors:** Elisabeth Roesch, Michael P.H. Stumpf

**Affiliations:** School of BioSciences and School of Mathematics and Statistics, University of Melbourne, Parkville VIC 3010, Australia; Centre for Integrative Systems Biology and Bioinformatics, Department of Life Sciences, Imperial College London, London SW7 2AZ, UK

## Abstract

Dynamical systems with intricate behaviour are all-pervasive in biology. Many of the most interesting biological processes indicate the presence of bifurcations, i.e. phenomena where a small change in a system parameter causes qualitatively different behaviour. Bifurcation theory has become a rich field of research in its own right and evaluating the bifurcation behaviour of a given dynamical system can be challenging. An even greater challenge, however, is to learn the bifurcation structure of dynamical systems from data, where the precise model structure is not known. Here we study one aspects of this problem: the practical implications that the presence of bifurcations has on our ability to infer model parameters and initial conditions from empirical data; we focus on the canonical co-dimension 1 bifurcations and provide a comprehensive analysis of how dynamics, and our ability to infer kinetic parameters are linked. The picture thus emerging is surprisingly nuanced and suggests that identification of the qualitative dynamics — the bifurcation diagram — should precede any attempt at inferring kinetic parameters.

## 1 Introduction

Modelling in the physical sciences often proceeds in a rigorous and disciplined manner: a small set of principles is sufficient to develop theoretical models of e.g. molecular dynamics, transport processes in solids, or transitions between different phases in condensed matter theory Anderson (1972). Symmetries then lead to conservation laws which can guide model development and greatly add in the interpretation of such models Neuenschwander (2011).

However, in many domains, including biology, modelling has to follow a different procedure May (2004): for example, if the basic symmetries are too far removed from the processes that we want to model, or the system is too complex (or indeed complicated) to model based on *first principles* Kirk et al. (2015). In these scenarios, we typically have to develop models based on domain expertise and subsequently compare them to available data using e.g. rigorous statistical model selection or model checking methods Kirk et al. (2013). There is, as a result, a vast literature on *reverse engineering* and *inverse problems* Moler and Loan (2003); Erguler and Stumpf (2011); Transtrum et al. (2010); Marbach et al. (2012); Transtrum et al. (2015); Herbach et al. (2017); Golightly and Wilkinson (2015). These sets of approaches allow us to develop models – typically iteratively – in light of available data and background information: we define the models, design more discriminatory experiments, make testable predictions about the behaviour of complex systems, and gain mechanistic insights into the inner workings of such systems. Reverse engineering and statistical model selection methods have found widespread use in many disciplines, ranging from engineering and biology to economics and the social sciences Transtrum et al. (2015).

Much of the literature in this area is focused on important systems where dynamics can be described or at least approximated in terms of linear differential equation models Transtrum et al. (2010); Tarantola (2013). For non-linear and stochastic dynamics, problems exacerbate very quickly. Here we address one aspect of reverse engineering that is of particular importance in many applied sciences. *Bifurcations* in non-linear dynamical systems result in a qualitative change in system behaviour; they are important in a host of biological processes, ranging from cell cycle control Tyson et al. (2003), cell fate decision making Moris et al. (2016); Bargaje et al. (2017), to ecological Sugihara et al. (2012) and epidemiological problems Jansen et al. (2003), and neurophysiology O’Donnell et al. (2017). In developmental biology, for example, we often seek to identify the stable stationary fixed points of an ODE system with distinct *cell states* Ferrell (2012); Moris et al. (2016). Equally, when considering an infectious disease we also distinguish different stationary points of the population dynamics with different meaning, e.g. disease is controlled vs. disease has spread. In modelling such systems we want to be able to capture the qualitative dynamics accurately. And this means that we have to be able to detect qualitative change points or bifurcations.

Here we focus on the simplest bifurcations in their simplest setting: scalar one -dimensional versions of four canonical types of bifurcations Jost (2006).

## 2 Methods

### Bifurcations

In this study, we investigate dynamical systems modelled by ordinary differential equations (ODE) that undergo *bifurcations*. Therefore, a system’s behaviour may differ qualitatively depending on small changes in parameters Jost (2006). Previous work has shown that the existence of bifurcations can profoundly affect our ability to infer parameters Kirk et al. (2008); Higham (2009) and here we extend this by considering and contrasting the canonical *co-dimension 1* bifurcations. These are bifurcations where the qualitative change is caused by the variation in a single parameter, which we refer to as the *bifurcation parameter α*. We look at them in their simplest normal form which are defined via the following ODEs:
*Saddle-node* bifurcation:

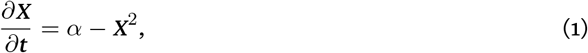

*Transcritical* bifurcation:

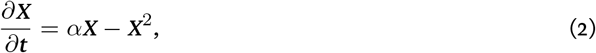

*Supercritical pitchfork* bifurcation:

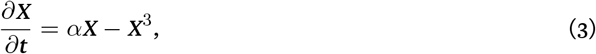

*Subcritical pitchfork* bifurcation:

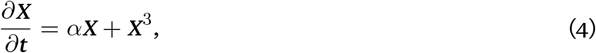

where *X* is the observable species and *t* is the time. Figure 1 depicts the bifurcation diagrams of these systems. In the first system a saddle-node bifurcation occurs at *α* = 0 (Fig. 1, 1. Saddle-node bifurcation). For *α* < 0 no fixed points exist and the system diverges whereas for *α* > 0 one stable and one unstable fixed points exist 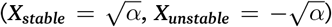. The second system undergoes a transcritical bifurcation (Fig. 1, 2. Transcritical bifurcation). Again, we observe the qualitative stability change at *α* = 0. For *α* < 0 and for *α* > 0 two fixed points exist; however, their locations vary (*α* < 0: *X*_*unstable*_ = −*α* and *X*_*stable*_ = 0, *α* > 0: *X*_*stable*_= *α* and *X*_*unstable*_ = 0).

**Figure 1:**
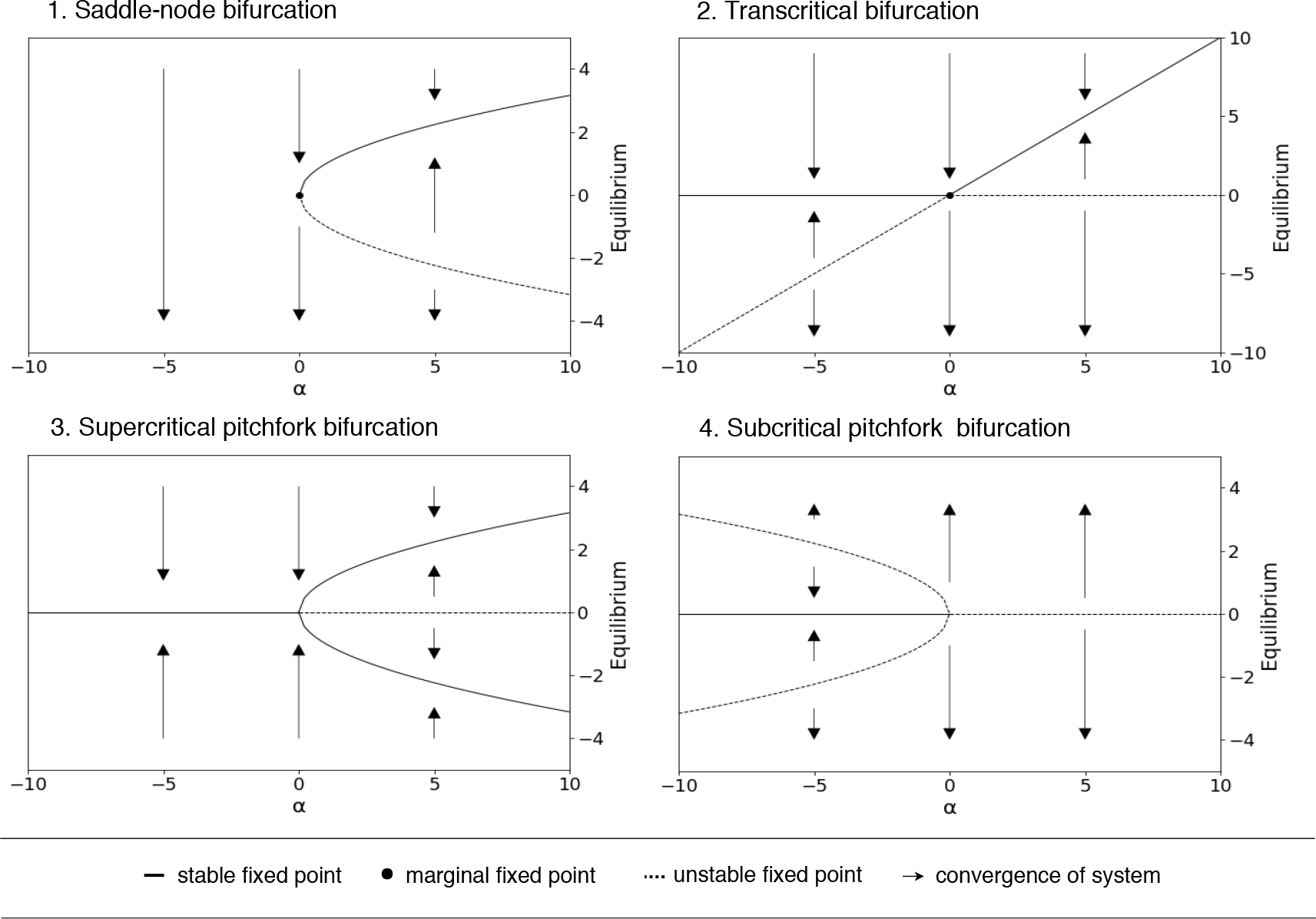
Bifurcation diagrams for the saddle–node, transcritical, and super/sub–critical pitchfork co–dimension 1 bifurcation. The diagrams visualize the global stability properties of dynamical systems(Eq. 1–4) depending on the bifurcation parameter α and the initial condition of the system IC. We see that due to small, smooth changes in the parameter space, the system’s behaviour changes in a qualitative manner as fixed points appear, disappear or change their stability properties. The graph represents fixed points which are stable (solid line) orunstable(dashedline) and arrows describe the convergence or divergence to /from a fixed point of a given system. If a fixed point is marginal it holds stable and unstable stability properties (dot).

The third and fourth system both exhibit pitchfork bifurcations representing the supercritical and subcritical case, respectively (Fig. 1, 3. Supercritical pitchfork bifurcation and 4. Subcritical pitchfork bifurcation). The system’s behaviour is symmetric along *X* = 0 for both types. However, they are mirror-inverted regarding the positioning of the fixed points but the stability properties of the fixed points are complementary (supercritical: *α* <= 0 : *X*_*stable*_= 0, *α* > 0 : 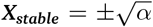 and *X*_*unstable*_= 0, and subcritical: *α* < = 0 : 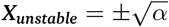 and *X*_*stable*_= 0, *α* > 0 : *X*_*unstable*_= 0).

### Likelihood

We investigate how the inference of parameters, but also of initial conditions, is affected by qualitative changes in these systems. In particular, we are trying to understand how the likelihood Cox (2006) over parameters and initial conditions is affected by the presence of a bifurcation. We follow the general approach of Kirk et al.Kirk et al. (2008). The log-likelihood Cox (2006) for such problems (assuming identically and independently normally distributed experimental noise) can be written, up to a constant proportionality factor, as

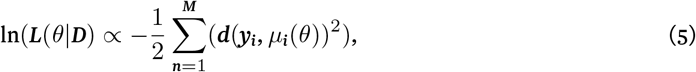

where *θ* = (*α*, *IC*) is the combined vector of rate parameter and initial condition; *D* = [*y*_1_, *y*_2_,…, *y*_*M*_] denotes the observations; and [*µ*_1_(*θ*), *µ*_2_(*θ*),…, *µ*_*M*_(*θ*)] represents the theoretical outputs for given *θ*; and where *d* is the Euclidean distance between the theoretical and observed data.

The value of *θ* for which Equation 5 becomes maximal is the *Maximum Likelihood estimate* (MLE) Cox(2006) and can be obtained numerically. Here, as we are interested in assessing how inference is affected by such changes, we calculate the likelihood over the *α*, *IC* plane. An intuitive explanation for the likelihood is given in Figure 2.

**Figure 2:**
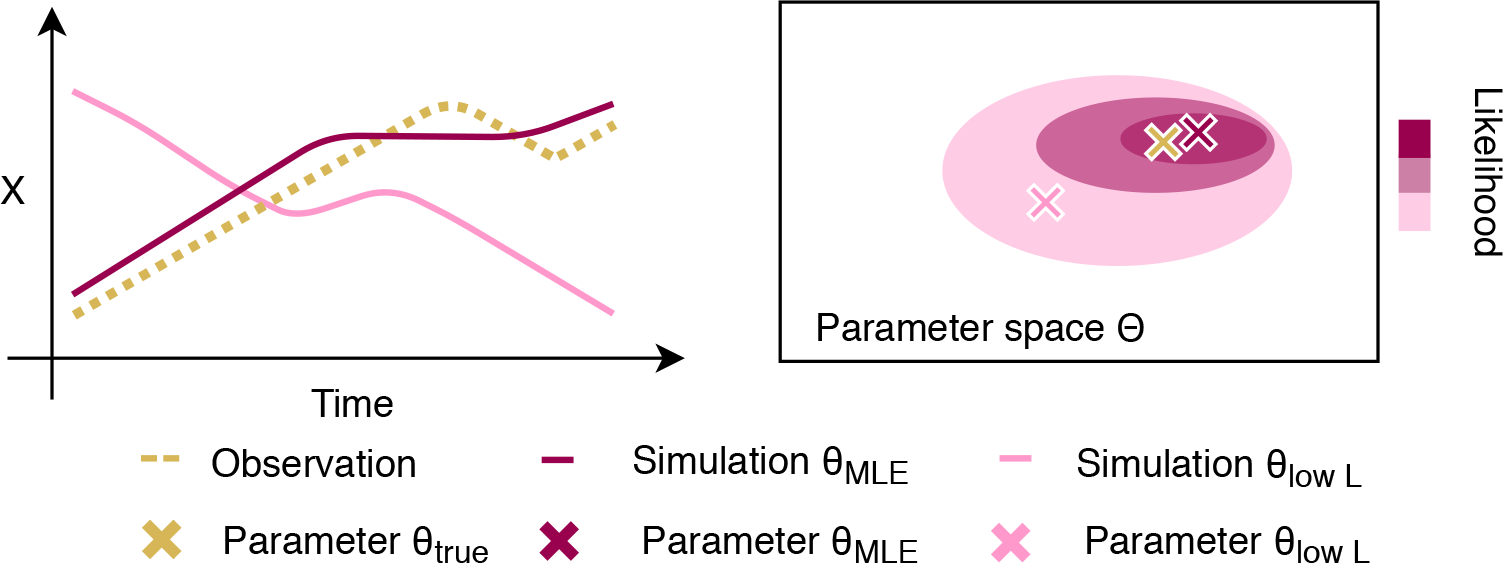
Intuition for the Likelihood function. The likelihood of a parameter combination θ expresses how well the simulation of θ describes the observation D. Therefore, the simulation of the model with a parameter combination with high likelihood such as θ_MLE_ represents the observed data well whereas the model simulation of the parameter combination θ_lowL_ differs significantly from the observation. Therefore, the true parameter combination θ_true_ is assumed to be closer to θ_MLE_ than θ_lowL_ in the parameter space Θ.

### Fisher Information

The MLE denotes the most likely value of the parameter given the observed data. It does not give an assessment of how much better is than a different value of the parameter. The Fisher informationCox (2006); Kirk et al. (2008); Komorowski et al. (2011) gives a more nuanced view of how reliably we can infer parameters, or how much certainty we should have in a given inferred parameter value: it is a measure of how quickly the likelihood changes around the MLE and needs to be understood as the curvature of the log-likelihood function, which is given by the second derivative (or the Hessian Matrix in the multivariate case),

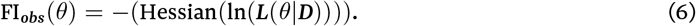

The entries, *H*_*ij*_ of the Hessian matrix, *H*, provide the curvatures,

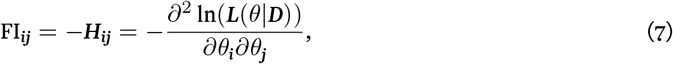

Intuitively, a more peaked log-likelihood surface around the MLE will have a higher Fisher information than a less peaked surface: this is because for the former, even a small shift away from the MLE will result in a substantial decrease of the log-likelihood, compared to the latter. The Fisher information quantifies this and is hence a measure of how well a parameter is inferred.

We summarise the Fisher Information matrix (as our preferred measure of inferability) by taking the trace of the matrix

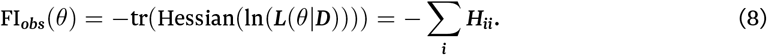

This value summarizes the information content that the observations *D* carry about about the bifurcation parameter *α* and the initial condition *IC*. An illustration is given in Figure 3.

**Figure 3:**
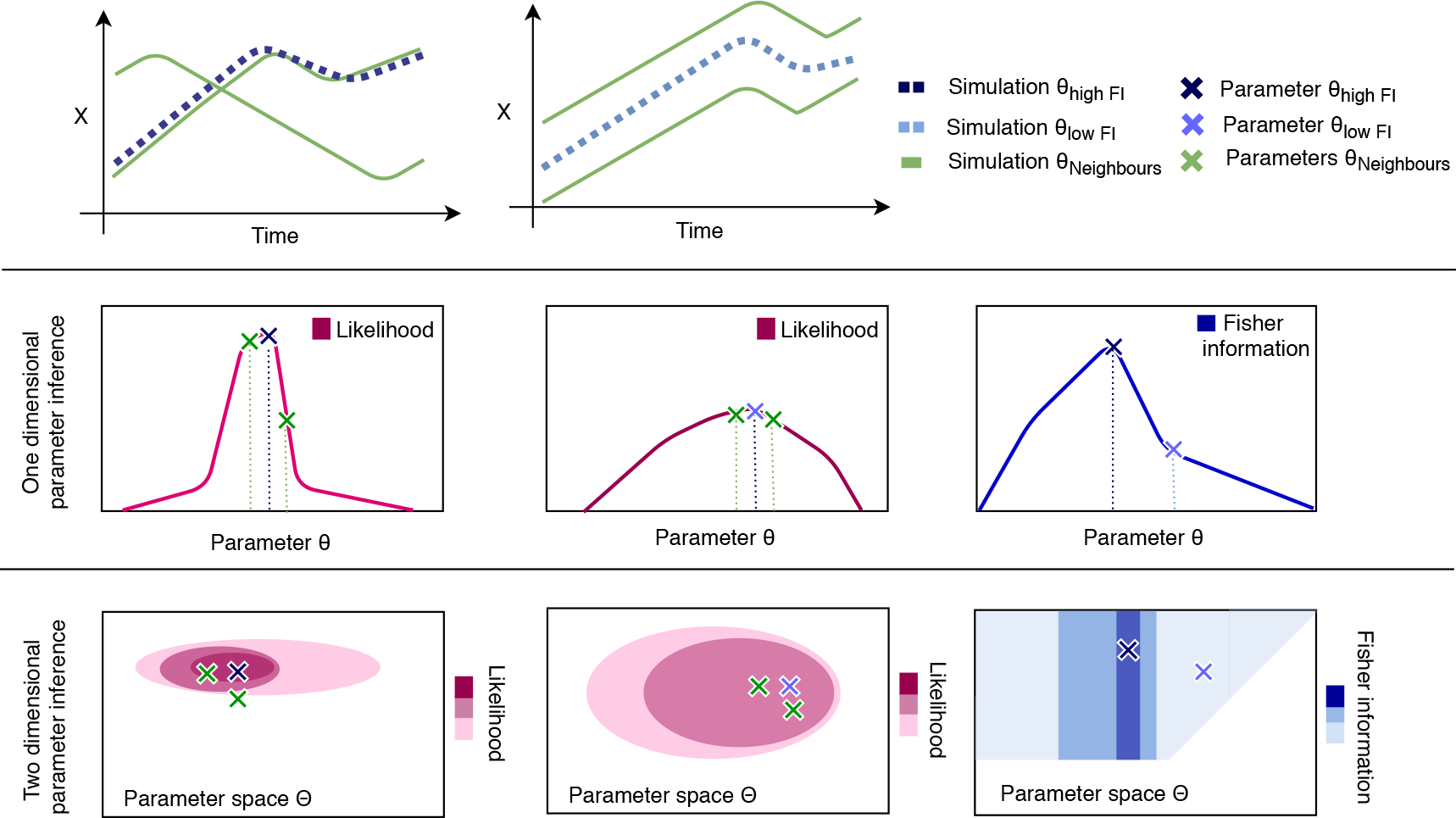
Intuition for the Fisher information. For a parameter combination with high Fisher information θ_highFI_ (the MLE of the likelihood function of θ_highFI_) neighbouring parameter samples may show significantly different likelihood values. If this is the case the distance of the simulations of the neighbouring parameters to the observation, and subsequently to the simulation of θ_highFI_, is large. Neighbouring parameter samples of θ_lowFI_ (the MLE of the likelihood function for θ_lowFI_), however, result in more similar values. Therefore the simulations of neighbouring parameter are equally distant to the observation and the simulation of θ_lowFI_. In this figure, we show the one dimensional case(e.g. θ = α) and the two dimensional case(e.g. theta = (α, IC). In the one dimensional case the blue line graph repesents the Fisher information and indicates for each value of θ the the observed curvatures of it’s on dimensional likelihood function. In the two dimensional case, the Fisher information is presented via a contour plot where intense/weak color indicates high/low Fisher information. The two dimensional likelihood surface of a parameter combination with high Fisher information θhighFI has a peaky shape (high curvature) whereas a parameter combination with low Fisher information θlowFI results in a broader shape (low curvature).

### Generating Simulated Data

Due to the imperfect nature of experimental data, measurements of the observation *D* are afflicted by noise. To assess the effects of such noise on parameter inference, we generate noisy data and investigate how the likelihood changes with increasing noise for the same parameter combination. The noise term follows a zero-centred *Gaussian* distribution 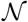 with standard deviation *σ*,

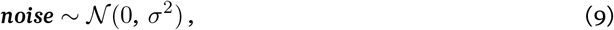

and noise levels are defined as low (*σ* = 0), moderate (*σ* = 0.1) and heavy (*σ* = 1.0); other noise models, e.g. log-normal are also possible, of course. Regarding the true parameter combination, we focused on *θ* = (0, 0.1) as we are particularly interested in the noise effect around the bifurcation area.

To better reflect many real-world problems we also consider the case of fewer data points, *M*, and the impact that this has for inference (which was also discussed at length in Kirk et al. (2008), and we consider very high (*M*=1000), high (*M*=500) and low (*M*=10) sampling rates.

## 3 Results

We first present the results for the saddle-node bifurcation, before summarizing the results for the three other bifurcations. Finally, we discuss the effects of sparse and noisy data on the parameter inference problem.

### Likelihood estimation around the saddle–node bifurcation

The parameters we wish to estimate are the bifurcation parameter, *α*, and the initial condition of the system, *IC*. In Figure 4, we select a group of exemplary parameter combinations covering the possible qualitative system behaviour and display the associated likelihood surfaces: no fixed point (*θ*_*a*_), single marginal fixed point (*θ*_*b*_)), and systems with one stable and one unstable fixed point (*θ*_*c*_-*θ*_*e*_).

**Figure 4:**
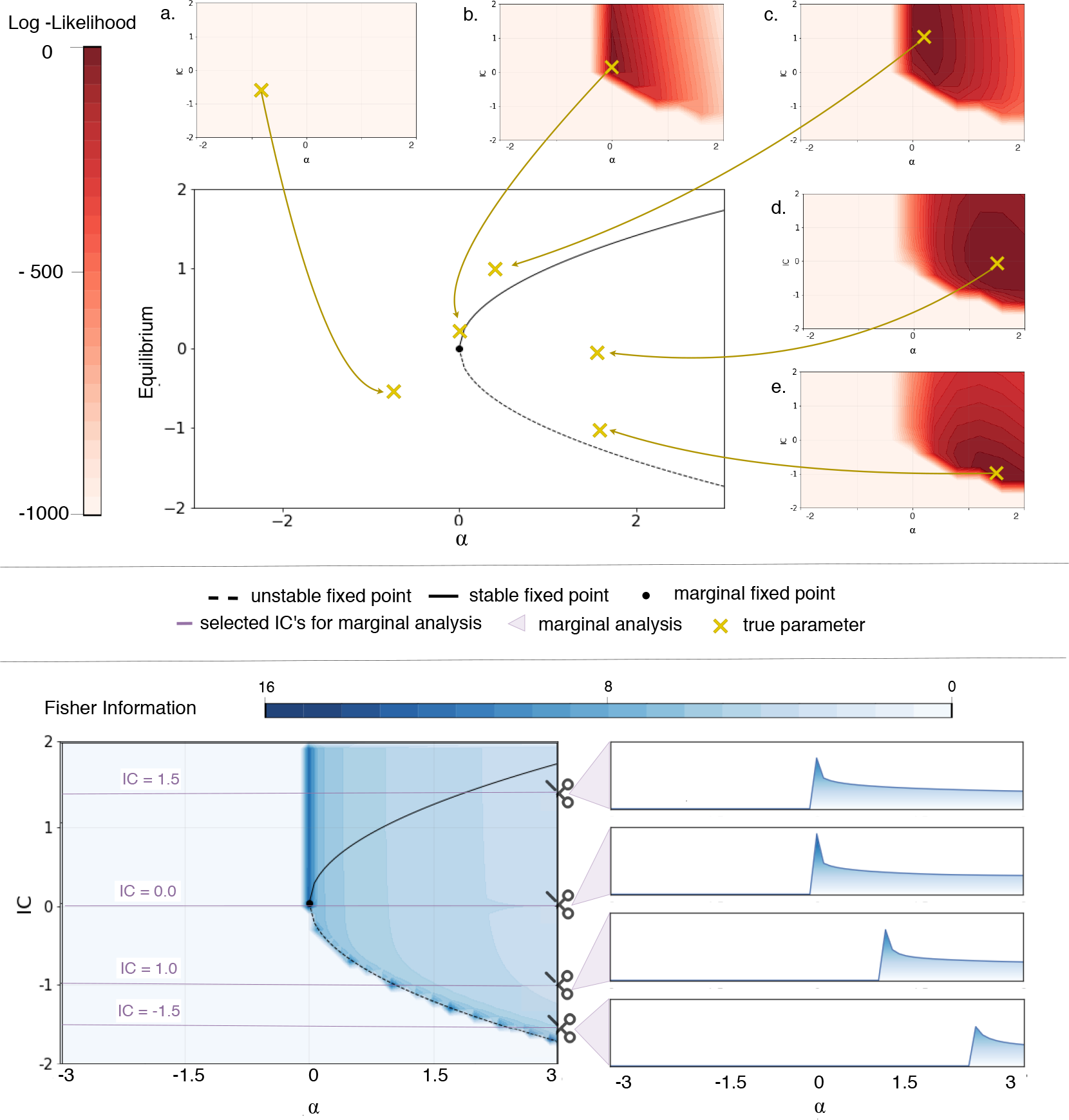
Quantifying information content around the saddle–node bifurcation via the log-likelihood function and Fisher information. In red, the log-likelihood surfaces of selected parameter combinations are depicted. They are surrounding the bifurcation diagram of the saddle–node bifurcation in order to set the selection of parameters in global stability context. Parameter combinations with (a) negative α give rise to diverging solutions. The likelihood function of a parameter combination close to the bifurcation event at α = 0 buton the non-negative side (b) shows a different behaviour and a pronounced peak can be observed. This is due to even minor changes of α causing the stability of the whole system to change. Therefore, parameter combinations can be inferred precisely in this area. Moving away from the bifurcation event but for α > 0 (c, d), we observe broader likelihood shapes. This means parameters can be inferred but the certainty decreases. Staying in the case where α > 0, but looking at parameter combinations closer to the unstable fixed point 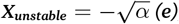 (e), the likelihood surfaces have more pronounced peaks again. This is due to the fact that the slightest change of IC to smaller values causes the system to diverge. In blue, the Fisher information is calculated for the whole parameter space. Each data point (α_x_, IC_y_) represents the observed Fisher information FI_obs_(θ_x,y_) where θ_x,y_ = (α_x_, IC_y_). This value summarizes the curvature of the log-likelihood for the parameter combination and we use it to measure the inferability of the parameters. Additionally, for five IC the margnialised Fisher information with increasing α is presented. For four IC the margnialised Fisher information with increasing α is presented.

The first parameter combination, *θ*_*a*_, represents combinations of negative *α* and arbitrary *IC*. As there are no fixed points in this region of the parameter space the systems do not converge to a stationary point but drifts off to −*∞*. Parameter and initial condition inference become essentially impossible as the log-likelihood decreases rapidly along these diverging system trajectories.

When *α* and *IC* become positive, the log-likelihood surface changes drastically. As an example, we show the likelihood surface of *θ*_*b*_, which is now very close to the bifurcation event at *α* = 0 and fulfills *α*, *IC* > = 0. We can identify a pronounced peak in the log-likelihood surface at the location of the true parameter combination, and accurate parameter inference becomes possible in this domain; sround *α* = 0, even slight changes in the parameter *α* can result in qualitatively different system dynamics. In the case of *θ_b_*, solutions with even infinitesimally smaller *α* will diverge, and with the likelihood surface we can rule such values out. A similar situation is observed for different initial conditions as *IC*s below the stationary point result in diverging solutions.

For the final category of parameter combinations we increase *α* further. Here, two fixed points exist, where one is stable and one is unstable. The parameter combination *θ*_*c*_ is located above the stable fixed point, whereas *θ*_*d*_ and *θ*_*e*_ are between the two fixed points. For all three parameter combinations we can estimate the true parameter from the peak of the likelihood surface which does indeed cover the true parameter combination. Also, like in the last scenario, the log-likelihood readily rules out any parameter combinations with negative *α* or *IC*. However, we observe that from *θ*_*b*_ to *θ*_*c*_ to *θ*_*d*_ the shape of log-likelihood surface broadens, i.e. inference becomes harder as we move away from the bifurcation event towards larger positive values of *α*. Initial conditions close to the unstable fixed points result in a more pronounced peak of the likelihood surface, as initial conditions below the unstable fixed points can be ruled out by the likelihood.

### Fisher information around the saddle–node bifurcation

Looking at the exemplar likelihood surfaces in parameter space gives us a good idea of the general ability to perform parameter inference around the bifurcation point. It allows us to investigate the overall shape of the log-likelihood and it enables us to assess how successful or straightforward parameter inference on different sides of the bifurcation will be. For example, in Figure 4, the log-likelihood surfaces of *θ*_*b*_ and *θ*_*d*_ both cover the true parameter. However, the quality of the two parameter estimates differs as the log-likelihood of *θ*_*b*_ is more peaked than the log-likelihood surface for *θ*_*d*_. If the (log)likelihood decays quickly around the MLE, we would put more certainty on this estimate, than if the surface is essentially flat in the vicinity of the maximum. This means the estimate for *θ*_*b*_ appears to be more informative, and therefore reliable, than the one for *θ*_*d*_. Formally, this is encapsulated by the Fisher Information (discussed above).

In Figure 4, the Fisher information *FI*_*obs*_ is calculated from the log-likelihood function of all parameter combinations across the parameter space. Each data point represents a summary of the curvature of the loglikelihood around the MLE for the specific parameter combination. As we can see, the existence and position of fixed points leave a profound mark in the Fisher information. Most importantly, the bifurcation event at *α* = 0 and the unstable fixed point at 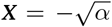 coincide with high-values of the Fisher information. We note that the stable fixed point at 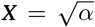 seems to have little bearing on the Fisher information and we do not observe it in the Fisher information at 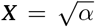. This is because all initial conditions 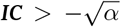 converge to it.

In many real applications we know the *IC* and only the kinetic (bifurcation) parameter is unknown and needs to be inferred. Therefore, we shift focus on the Fisher information of the parameters but for known, specific *IC*s (Fig. 4). We chose a collection of four *IC*s capturing all general marginal structures found in the saddle– node bifurcation.

First, we set *IC* to 1.5. In this case we observe the highest Fisher information at the bifurcation event where *α* = 0; for positive *α*, the Fisher information decrease when moving away from the bifurcation event. Marginalizing the Fisher information at *IC* = 0 results in a similar general shape, however the peak of the Fisher information at 0 is even more pronounced. For *IC* = −1.0 we find again the same general structure, but the maximum’s location is now shifted away from the bifurcation event to the unstable fixed point at 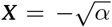 and the same is true for *IC* = −1.5.

### Fisher information around the other bifurcations

In order to outline the results of the transcritical, supercritical pitchfork and subcritical pitchfork bifurcation we show the Fisher information plots in Figure 5. Log-likelihood surfaces for selected parameter combinations are shown in the appendix for the transcritical (Figure 7), supercritical pitchfork (Figure 8) and subcritical pitchfork bifurcation (Figure 9).

**Figure 5:**
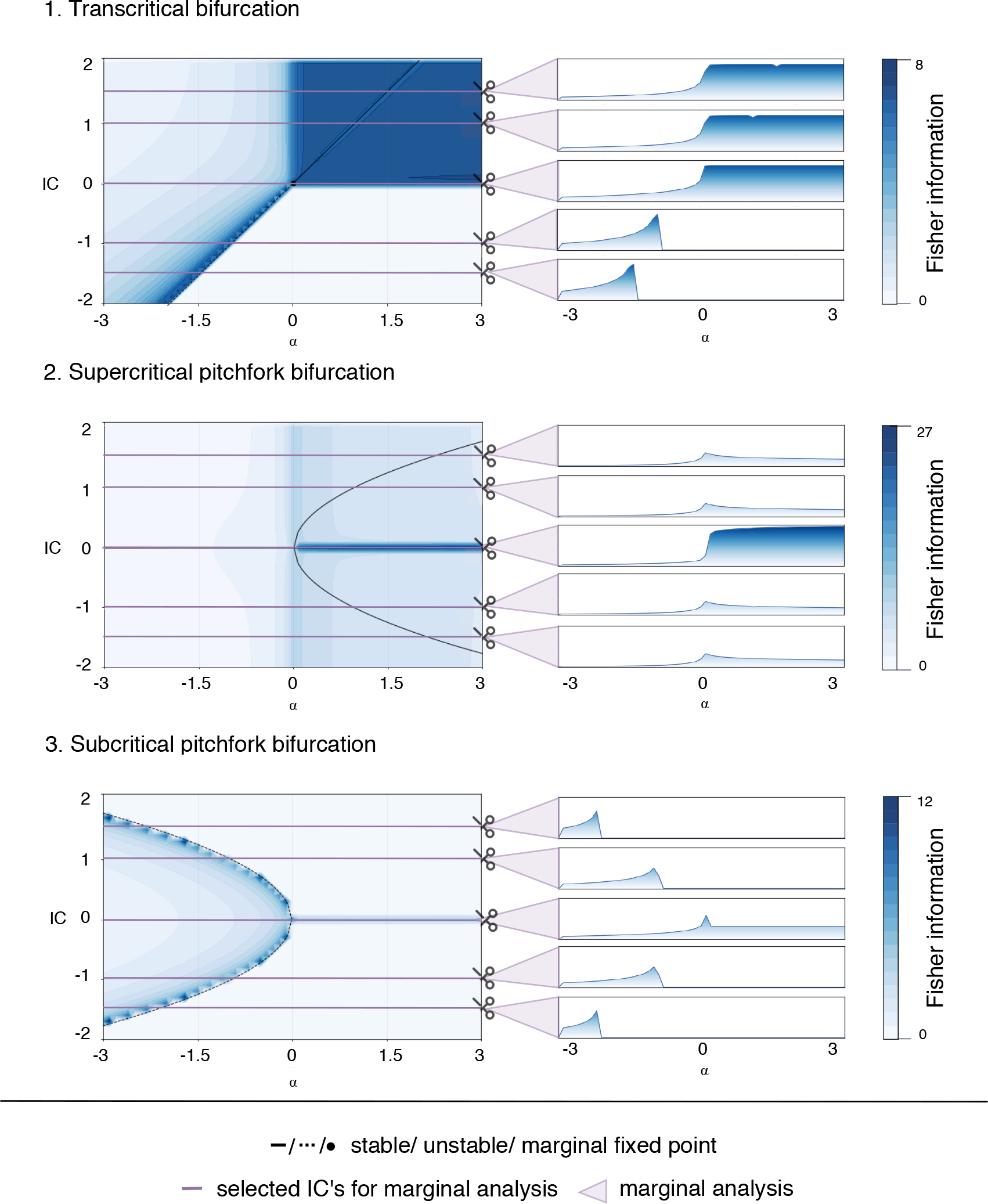
Quantifying information content around the transcritical, supercritical and subcritical pitchfork bifurcation via Fisher information. For each bifurcation, the Fisher information is calculated for the whole parameter space. Each data point (α_x_, IC_y_) represents the observed Fisher information FI_obs_(θ_x,y_) where θ_x,y_ = (α_x_, IC_y_). This value summarizes the curvature of the log-likelihood for the parameter combination and we use it to measure the inferability of the parameters. In order to emphasize the connection of qualitative stability properties and the inferability, the bifurcation diagrams are added. Additionally, for five IC the margnialised Fisher information with increasing α is presented. For all three bifurcations we observe enriched Fisher information at the bifurcation event (transcritical: α = 0, supercritical pitchfork: α = 0, subcritical pitchfork: α = 0) and along unstable fixed points (transcritical: for α < 0 at X_unstable_ = α and for α > 0 at X_unstable_ = 0, supercritical pitchfork: α > 0 at X_unstable_ = 0, subcritical pitchfork: α < 0 at 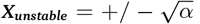 and for α > 0 at X_unstable_ = 0). Equivalently to the saddle-node bifurcation and due to visualisation purposes, the Fisher information is not calculatable and therefor set to 0 if the parameter combination causes the system to diverge.

**Figure 6:**
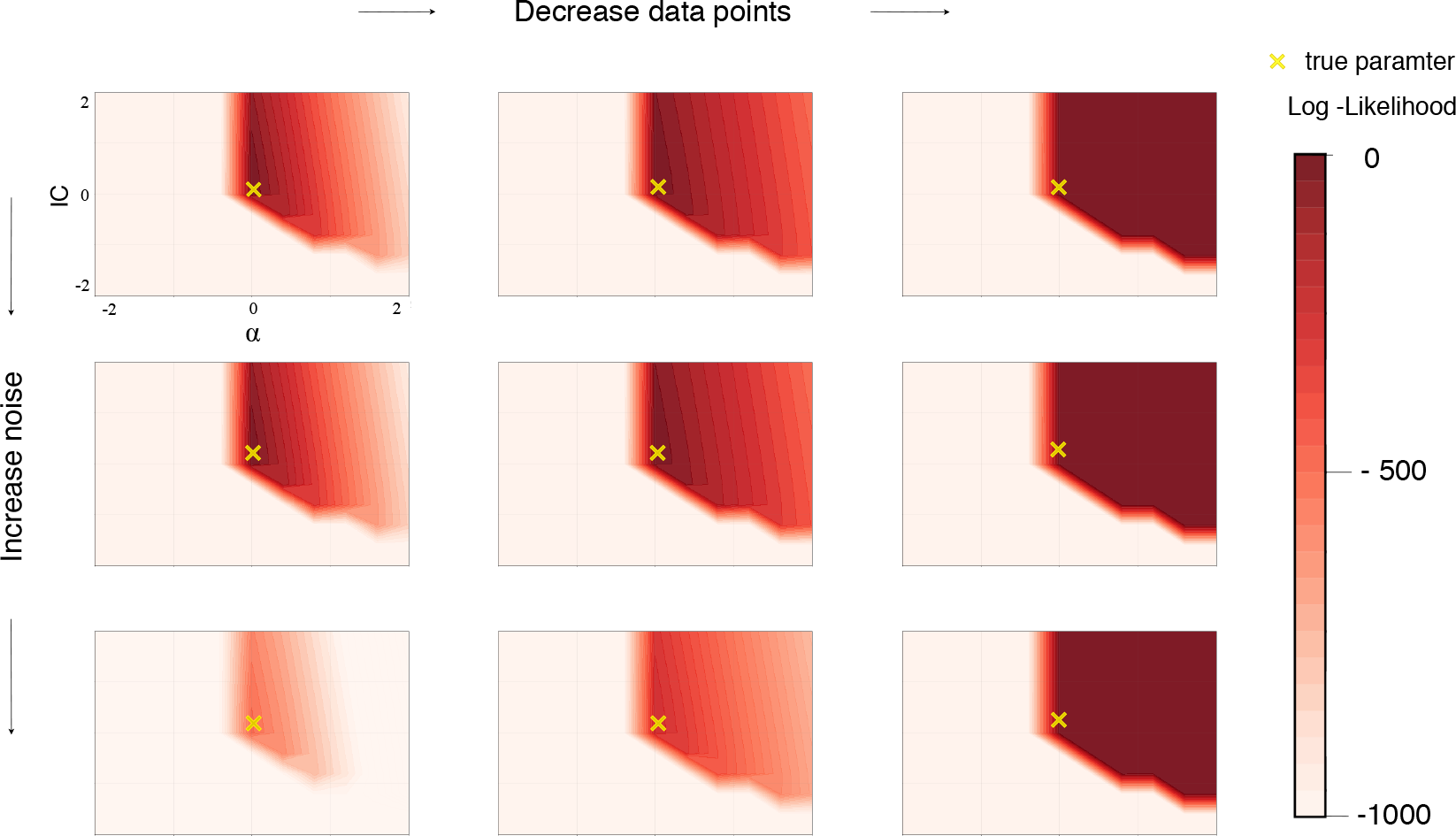
Effect of noisy and sparse data on the shape of the log-likelihood function for the Saddle-node bifurcation. From top to bottom we increase the noise (σ: 0, 0.1, 1.0) and from left to right we decrease the number of observations (M = 1000, 500, 10). The true parameter is always kept the same. We observe that the number of parameter combinations.

**Figure 7:**
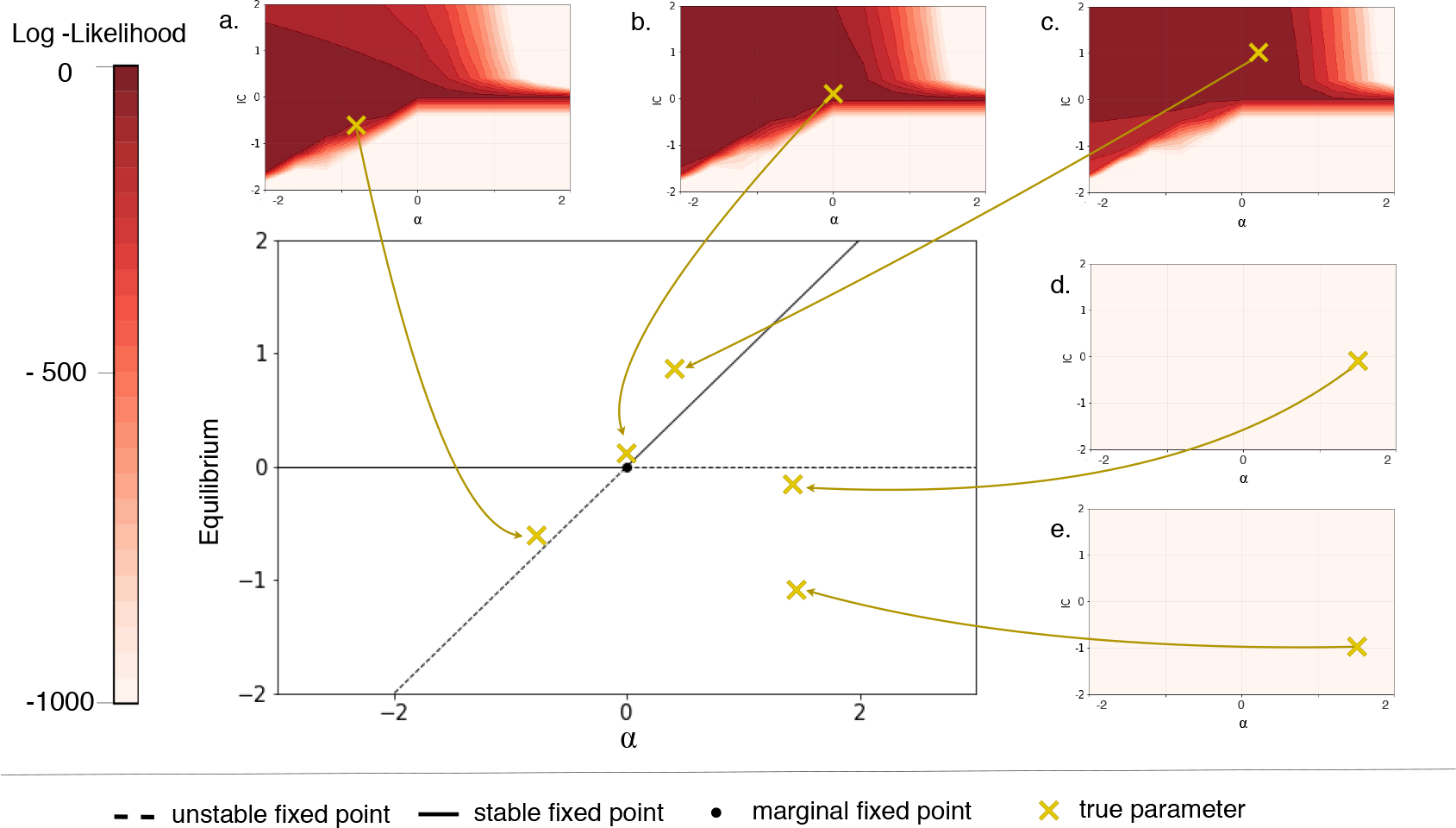
Quantifying information content around the transcritical bifurcation via the Log-likelihood. Selected parameter combinations are highlighted in the bifurcation diagram in gold and their log-likelihood is shown in red. Parameter combinations with negative α and IC > α (a), α = 0 and IC > 0 (b), or positive α and IC > 0 (c), can be inferred. However, parameter combinations below the unstable fixed points such (d) and (e), cannot be inferred.

**Figure 8:**
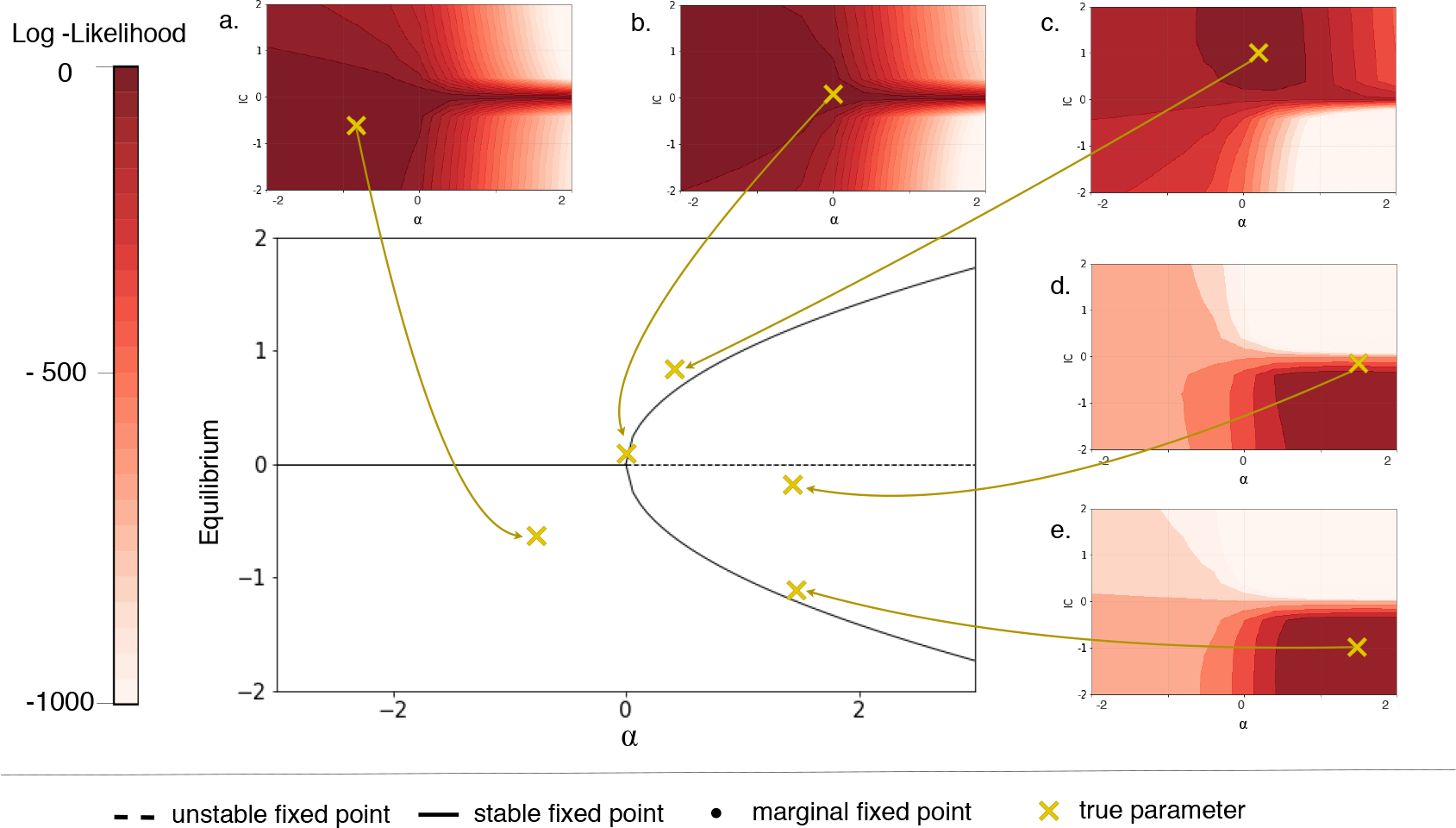
Quantifying information content around the supercritical pitchfork bifurcation via the Log-likelihood. Selected parameter combinations are highlighted in the bifurcation diagram in gold and their log-likelihood is shown in red. For this bifurcation, parameter inference can be performed in every area of the parameter space.

**Figure 9:**
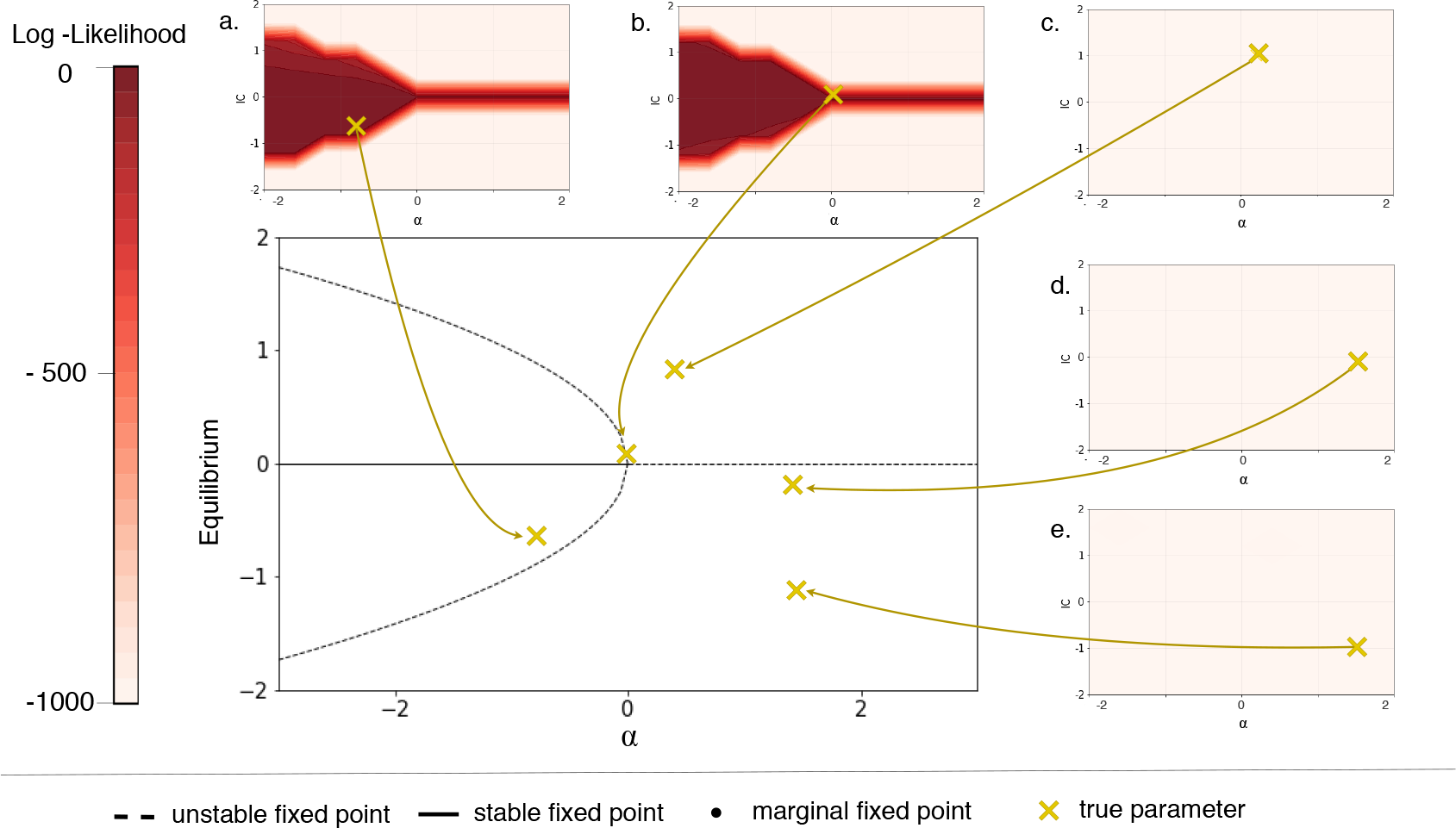
Quantifying information content around the subcritical pitchfork bifurcation via the Log-likelihood. Selected parameter combinations are highlighted in the bifurcation diagram in gold and their log-likelihood is shown in red. Due to the unstable fixed points, parameter inference is the hardest for this bifurcation. We show, only parameter combinations with negative α and 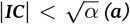 (a), or, α = 0 and IC close to zero (b) can be inferred. Note, that the true system for (b) is actually diverging but due to slow divergence compared to the chosen simulated time span, the inference procedure works.

#### Transcritical bifurcation

The transcritical bifurcation varies considerably from the saddle-node bifurcation as fixed points exists for every value of the bifurcation parameter *α*. However, we still observe major differences in the parameter inferability for *α* < 0 and *α* > 0. For *α* < 0, high Fisher information and therefore peaked likelihood surfaces are found for parameter combinations close to, but above, the unstable fixed point 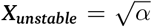. Certainty of the inference (as measured by the Fisher information) decrease as we move further away from the fixed point. In the case of *α* > 0 and positive *IC*s parameter combinations can be inferred with high certainty. The whole quadrant shows consistently high Fisher information. Interestingly, this is the only bifurcation in which the stable fixed point is clearly visible in the Fisher information (for negative *α*).

#### Superscritical pitchfork bifurcation

For the supercritical pitchfork bifurcation, we also find at least one fixed point for every *α*. Similarly to the transcritical bifurcation we still observe variation in the inferability for positive and negative values of *α*. However, we observe symmetry along the *α*-axis. This is in line with our expectations, as the bifurcation diagram is symmetric, too. The highest certainty is associated with the unstable fixed point *X*_*stable*_ = 0. It decreases when moving away from the unstable fixed point but generally for *α*> 0 the likelihood surfaces are more peaked then for negative *α*. It is worth mentioning that for this bifurcation, the predictions for negative *α* are still fairly certain even though they are colored similarly to less certain predictors of other bifurcations but this is only due to the varying color bars (see supplementary material for the likelihood surfaces).

#### Subscritical pitchfork bifurcation

The subcritical bifurcation is the bifurcation with the most diverging dynamics of the ones considered here: For many *IC* and *α* combinations the trajectories will not converge to a stable fixed point. We are still able to identify the unstable fixed points very clearly from the Fisher information. Moving away from the unstable fixed point decreases the certainty of predictions.

### Noisy and sparse data

In the discussion above we have focused on idealised data that is noiseless and plentiful. Relaxing these ideal conditions does, reassuringly, not alter our findings qualitatively. as shown in Figure 6, making the data sparser and adding noise still results in likelihood and Fisher information behaviours that recapitulate what we have described above.

Thus, like in the case of the Hopf-bifurcation previously studied by Kirk et al.Kirk et al. (2008), we find that the qualitative hallmarks of the dynamics induced by the bifurcations persist in less than ideal data.

## 4 Conclusion

We have demonstrated that the structure of a dynamical system, and the qualitative manifestations of system dynamics affect our ability to learn and infer dynamical parameters as well as initial conditions. Here we have only considered the simplest examples of bifurcating systems, but these still showed diverse and subtle behaviour. Most importantly, the *global* dynamics can affect profoundly our ability to make *local* parameter inferences. Our discussion of results for more realistic (sparse and noisy) data suggests, that in practice we should be able to detect the hallmarks of qualitatively different dynamics in parameter inference studies. But our analysis also suggests that some parameters will be systematically more difficult to infer than others, depending on the type of bifurcation under consideration. If a parameter is harder to infer (as e.g. measured by the Fisher information) this means that parameters in its vicinity will result in very similar system behaviour.

In real-world dynamical systems, e.g. in population and systems biology, we often expect several of these bifurcation types to be present Moris et al. (2016). Disentangling the interplay between global qualitative dynamics and our ability to infer parameter (or parameter combinations) will be more complicated. Some simple guidelines or heuristics, however, emerge from the present work, and these can be used as guidelines in more complicated scenarios: first, and perhaps most importantly, capturing the qualitative dynamical regime(s) qualitatively using mechanistic modelling will allow us to triage different hypotheses quickly based on even modest qualitative data; and second, the realisation that exploring different initial conditions can greatly help in parameter inference; thus experimental design will be crucial in practice as has previously been demonstrated Liepe et al. (2013); Silk et al. (2014).

## Acknowledgments

We gratefully acknowledge discussions with members of the Theoretical Systems Biology group in both hemispheres. MPHS has had a long-standing interest in this problem and has initially discussed this with Erika Cule and the late Jaroslav Stark, who have both helped to shape and refine the questions addressed here.

## 5 Appendix

Here we present and summarise graphically the results for the transcritical, super and sub-cirtical pitchfork bifurcations.

## References

P Anderson. More Is Different. Science, pages 393–396, August 1972.

Rhishikesh Bargaje, Kalliopi Trachana, Martin N Shelton, Christopher S McGinnis, Joseph X Zhou, Cora Chadick, Savannah Cook, Christopher Cavanaugh, Sui Huang, and Leroy Hood. Cell population structure prior to bifurcation predicts efficiency of directed differentiation in human induced pluripotent cells. Proceedings of the National Academy of Sciences, 114(9):2271–2276, February 2017.

D.R. Cox. Principles Of Statistical Inference. Cambridge University Press, 2006. ISBN 9780521685672.

Kamil Erguler and Michael P. H. Stumpf. Practical limits for reverse engineering of dynamical systems: a statistical analysis of sensitivity and parameter inferability in systems biology models. Molecular BioSystems, 7:1593–1602, 2011. doi: 10.1039/C0MB00107D. URL http://dx.doi.org/10.1039/C0MB00107D.

James E Ferrell, Jr. Bistability, bifurcations, and Waddington’s epigenetic landscape. Current Biology, 22(11): R458–66, Jun 2012. doi: 10.1016/j.cub.2012.03.045.

A Golightly and Darren J Wilkinson. Bayesian inference for Markov jump processes with informative observations. Statistical applications in genetics and molecular biology, 14(2):169–188, April 2015.

Ulysse Herbach, Arnaud Bonnaffoux, Thibault Espinasse, and Olivier Gandrillon. Inferring gene regulatory networks from single-cell data: a mechanistic approach. BMC Systems Biology, 11(1):105, November 2017.

Catherine F Higham. Bifurcation analysis informs Bayesian inference in the Hes1 feedback loop. Bmc Systems Biology, 3(1):12, January 2009.

Vincent A A Jansen, N Stollenwerk, H J Jensen, M E Ramsay, W J Edmunds, and C J Rhodes. Measles Outbreaks in a Population with Declining Vaccine Uptake. Science, 301(5634):804–804, August 2003.

Jürgen Jost. Dynamical Systems. Examples of Complex Behaviour. Springer, March 2006.

P Kirk, P D W Kirk, T Toni, Michael P H Stumpf, and Michael P H Stumpf. Parameter inference for biochemical systems that undergo a Hopf bifurcation. Biophysical journal, 95(2):540–549, July 2008.

P Kirk, Thomas W Thorne, and Michael P H Stumpf. Model selection in systems and synthetic biology. Current Opinion in Biotechnology, 24(4):767–774, August 2013.

P D W Kirk, Ann C Babtie, and Michael P H Stumpf. Systems biology (un)certainties. Science, 350(6259): 386–388, October 2015.

Michal Komorowski, Maria J Costa, David A Rand, and Michael P H Stumpf. Sensitivity, robustness, and identifiability in stochastic chemical kinetics models. Proceedings of the National Academy of Sciences, 108(21):8645–50, May 2011. doi: 10.1073/pnas.1015814108.

Juliane Liepe, S Filippi, M Komorowski, and Michael P H Stumpf. Maximizing the information content of experiments in systems biology. PLoS Computational Biology, 9(1):e1002888, 2013.

Daniel Marbach, James C Costello, Robert Küffner, Nicole M Vega, Robert J Prill, Diogo M Camacho, Kyle R Allison, DREAM5 Consortium, Manolis Kellis, James J Collins, and GUSTAVO Stolovitzky. Wisdom of crowds for robust gene network inference. 9(8):796–804, August 2012.

Robert M May. Uses and abuses of mathematics in biology. Science, 303(5659):790–793, February 2004.

Cleve Moler and Charles Van Loan. Nineteen dubious ways to compute the exponential of a matrix, twenty-five years later. SIAM Review, 45(1):3–49, 2003. doi: 10.1137/S00361445024180. URL http://dx.doi.org/10.1137/S00361445024180.

Naomi Moris, Cristina Pina, and Alfonso Martinez Arias. Transition states and cell fate decisions in epigenetic landscapes. Nature reviews. Genetics, 17(11):693–703, September 2016.

Dwight E Neuenschwander. Emmy Noether’s Wonderful Theorem. JHU Press, March 2011.

Cian O’Donnell, J Tiago Gonçalves, Carlos Portera-Cailliau, and Terrence J Sejnowski. Beyond excitation/inhibition imbalance in multidimensional models of neural circuit changes in brain disorders. eLife, 6:3116, October 2017.

Daniel Silk, P Kirk, Chris P Barnes, T Toni, and Michael P H Stumpf. Model selection in systems biology depends on experimental design. PLoS Computational Biology, 10(6):e1003650, June 2014.

George Sugihara, Robert M May, Hao Ye, Chih-hao Hsieh, Ethan Deyle, Michael Fogarty, and Stephan Munch. Detecting Causality in Complex Ecosystems. Science, 338(6106):496–500, October 2012.

A Tarantola. Inverse Problem Theory. Methods for Data Fitting and Model Parameter Estimation. Elsevier, October 2013.

Mark K Transtrum, Benjamin B Machta, and James P Sethna. Why are nonlinear fits to data so challenging? Physical Review Letters, 104(6):060201, February 2010. doi: 10.1103/PhysRevLett.104.060201. URL https://link.aps.org/doi/10.1103/PhysRevLett.104.060201.

Mark K Transtrum, Benjamin B Machta, Kevin S Brown, Bryan C Daniels, Christopher R Myers, and James P Sethna. Perspective: Sloppiness and emergent theories in physics, biology, and beyond. The Journal of chemical physics, 143(1):010901, July 2015.

John J Tyson, Katherine C Chen, and Béla Novák. Sniffers, buzzers, toggles and blinkers: dynamics of regulatory and signaling pathways in the cell. Current opinion in cell biology, 15(2):221–231, April 2003.

